# Assessing the pathogenicity, penetrance and expressivity of putative disease-causing variants in a population setting

**DOI:** 10.1101/407981

**Authors:** Caroline F. Wright, Ben West, Marcus Tuke, Samuel E. Jones, Kashyap Patel, Thomas W. Laver, R. N. Beaumont, Jessica Tyrrell, Andrew R. Wood, Timothy M. Frayling, Andrew T. Hattersley, Michael N. Weedon

## Abstract

Over 100,000 genetic variants are classified as disease-causing in public databases. However, the true penetrance of many of these rare alleles is uncertain and may be over-estimated by clinical ascertainment. As more people undergo genome sequencing there is an increasing need to assess the true penetrance of alleles. Until recently, this was not possible in a population-based setting. Here, we use data from 388,714 UK Biobank (UKB) participants of European ancestry to assess the pathogenicity and penetrance of putatively clinically important rare variants.

Although rare variants are harder to genotype accurately than common variants, we were able to classify 1,244 of 4,585 (27%) putatively clinically relevant rare variants genotyped on the UKB microarray as high-quality. We defined “rare” as variants with a minor allele frequency of <0.01, and “clinically relevant” as variants that were either classified as pathogenic/likely pathogenic in ClinVar or are in genes known to cause two specific monogenic diseases in which we have some expertise: Maturity-Onset Diabetes of the Young (MODY) and severe developmental disorders (DD). We assessed the penetrance and pathogenicity of these high-quality variants by testing their association with 401 clinically-relevant traits available in UKB.

We identified 27 putatively clinically relevant rare variants associated with a UKB trait but that exhibited reduced penetrance or variable expressivity compared with their associated disease. For example, the P415A *PER3* variant that has been reported to cause familial advanced sleep phase syndrome is present at 0.5% frequency in the population and associated with an odds ratio of 1.38 for being a morning person (*P*=2×10^-18^). We also observed novel associations with relevant traits for heterozygous carriers of some rare recessive conditions, e.g. heterozygous carriers of the R799W *ERCC4* variant that causes Xeroderma pigmentosum were more susceptible to sunburn (one extra sunburn episode reported, *P*=2×10^-8^). Within our two disease subsets, we were able to refine the penetrance estimate for the R114W *HNF4A* variant in diabetes (only ~10% by age 40yrs) and refute the previous disease-association of *RNF135* in developmental disorders.

In conclusion, this study shows that very large population-based studies will help refine the penetrance estimates of rare variants. This information will be important for anyone receiving information about their health based on putatively pathogenic variants.

## INTRODUCTION

One of the ongoing challenges in genetic medicine is that of variant interpretation. Many variants and genes have been erroneously associated with disease as a result of problems with study design, including ascertainment bias and inadequate cohort size^1-3^, as well as biological phenomena such as genetic heterogeneity, reduced penetrance, variable expressivity, composite phenotypes, pleiotropy and epistasis^4-13^. These issues have resulted in ambiguity over how to interpret clinically-ascertained variants found in individuals with no known family history or symptoms of the disease^14^. Although there has traditionally been a division between rare disease genetics (studied in small disease cohorts and individual high-risk families) and common disease genetics (studied in large disease cohorts and population biobanks), in reality there is likely to be a continuum of causality that exists for many human disorders^15^. Fortunately, rare and common disease studies suffer from opposing ascertainment biases. Clinically-ascertained cohorts are enriched for individuals with a specific clinical presentation, and will therefore tend to over-estimate the penetrance of any disease-causing variants identified^16^. In contrast, population cohorts tend to be enriched for healthy individuals (so-called “healthy volunteer” selection bias) who have both the time and ability to volunteer for a study^17,18^, and will therefore tend to under-estimate penetrance. Population cohorts with high-resolution genetic and clinical data are therefore invaluable for establishing minimum penetrance estimates, exploring variable expressivity and challenging pathogenicity assertions made in the clinical arena.

Several studies have already started to bridge this gap by using population data to evaluate rare disease-causing variants^19,20^, refine penetrance estimates^21^ and refute reportedly pathogenic variants^22,23^. These previous studies were mostly limited to a very specific set of variants (e.g. protein truncating), or one particular disease, or were too small to statistically test phenotypic penetrance. With its wealth of linked phenotypic and clinical information on >450,000 genotyped individuals, UK Biobank (UKB)^24^ offers a powerful dataset in which to systematically evaluate the pathogenicity, penetrance, and expressivity of clinically important variants in the population. However, differences in the technologies used to assay genetic variation can hinder these analyses. A particular concern is the use of genotyping arrays (such as those currently used by UKB)^25^, which have been designed primarily to assay common variation. In contrast, rare single nucleotide variants (SNVs) and small insertions/deletions (indels) have typically been detected through sequencing assays^26^. A method is therefore needed to select well-genotyped rare variants in UKB, which can then be used to address biological and clinical questions.

Here we describe a systematic method for evaluating the analytical validity of rare variant genotyping data from the UKB arrays, investigate the relationship between data quality and minor allele frequency (MAF), and evaluate the association of a subset of clinically-interesting, well-genotyped coding variants with relevant phenotypes in UKB. We focus on variants in ClinVar that have been classified as “pathogenic/likely pathogenic” by at least one submitter^27^, as well as variants in genes known to cause two specific monogenic diseases in which we have some expertise: maturity-onset diabetes of the young (MODY) and developmental disorders (DD).

## METHODS

### UKB cohort

UKB recruited over 500,000 individuals aged 37-73 years between 2006-2010 from across the UK. Participants provided a range of information via questionnaires and interviews (e.g. demographics, health status, lifestyle) and anthropometric measurements, blood pressure readings, blood, urine and saliva samples were taken for future analysis. Genotypes for single nucleotide variants (SNVs) and insertions/deletions (indels) were generated from the Affymetrix Axiom UKB array (~450,000 individuals) and the UKBiLEVE array (~50,000 individuals) in 106 batches of ~4,700 samples. This dataset underwent extensive central quality control (http://UKB.ctsu.ox.ac.uk)^25^. We limited our analysis to 388,714 QC-passed white Europeans.

### Variant prioritisation

Variants were annotated using Annovar^28^ and MAFs were calculated using PLINK^29^. To prioritise variants of potential clinical importance, we selected those with at least one classification of pathogenicity (pathogenic or likely pathogenic) in the ClinVar database (https://www.ncbi.nlm.nih.gov/clinvar/)^27^, including those with conflicting classifications. In addition, irrespective of their presence in ClinVar, we selected predicted protein truncating variants (PTV; stopgain SNVs and frameshift indels) and known pathogenic functional variants (nonsynonymous SNVs and inframe indels) in genes known to cause MODY^30,31^ (https://www.diabetesgenes.org) and dominant DD^32,33^ (https://www.ebi.ac.uk/gene2phenotype) for detailed evaluation. These diseases and genes were selected due to our own prior experience, the availability of well-curated gene lists that include mode of inheritance and mechanism of action, and the different priors associated with finding diabetes (a common disease) and severe DD (a rare disease) in UKB. We excluded common variants (MAF>0.01), as these have already been thoroughly investigated through genome-wide association studies^34,35^, and further refined the list of variants to include only those where the Hardy–Weinberg equilibrium (HWE) *P*>0.05 and the proportion of missing genotypes across all samples <0.01 (n=4,585).

### Assessing analytical validity

To assess the analytical validity of these variants, we used Evoker Lite (https://github.com/dlrice/evoker-lite) to generate cluster plots of intensities, and combined data from all the batches into one plot for each variant. Cluster plots were manually assessed and ranked in quality from 1-5, where: 1=poor quality, no discernible separate clusters; 2=poor quality, no discernible separate clusters but noisy data; 3=unclear/uncertain; 4=good quality, clearly separable clusters but noisy data; and 5=good quality, clear separation between clusters (**Supplementary Figure 1**). In an initial subset of 750 variants that was independently evaluated by two scientists (**Supplementary Figure 2**), correlation between the two independent scorers was high (R^2^=0.8), and there was a 95% agreement in low quality (score=1 or 2) versus high quality (score=4 or 5) variants. All remaining variants of interest were evaluated by one scientist, and those with high quality scores were checked by the second scientist. Only variants with an average score of >4 were retained for further analysis. For all 1,244 high-quality variants, we assessed whether the rare genotype calls were unusually distributed across the 107 genotype batches. None of the rare genotypes calls at these variants were entirely due to calls from a single batch. Across the 1,244 variants, the highest proportion of rare genotype calls in a single batch was 4 from a total of 13 for Affx-89007317. A plot of total allele count for each variant against maximum allele count across each individual batch demonstrated a linear association with no clear outlying variants.

### Assessing clinical relevance

We ran a phenome-wide association for all of our 1,244 high-quality rare variants against a curated list of 401 clinically-relevant traits in UKB (**Supplementary Table 1**) in 388,714 QC-passed white Europeans using PLINK^29^, and those with a Bonferroni-corrected p <1×10^-7^ (0.05/(401*1244)) were prioritised for detailed evaluation. For continuous traits, we used linear regression adjusting for age, sex (unless a sex-specific trait), centre, chip and ten ancestry principal components. For binary traits, we used Fisher’s exact test as the primary association method and performed logistic regression adjusting for the same covariates as for continuous traits as a sensitivity analysis. We excluded variants now considered by recent reclassifications in ClinVar to be benign. To assess the potential clinical implications of high-quality rare variants, we compared the UKB traits with the clinical presentation of the disease for each gene, and the evidence supporting the assertion of pathogenicity of the variant using ClinVar^27^, DECIPHER^36^ and OMIM^37^. For high-quality rare variants in MODY genes and PTVs in DD genes, we had no p-value cut-off for investigating diabetes and developmental traits (cognitive function, educational attainment, body mass index, height, hearing and albumin creatinine ratios). Conditional analysis of the most-associated regional variant (1Mb window) from each trait led us to remove one trait-variant association that was explained by linkage disequilibrium with a common causal variant.

## RESULTS

### Variants below 0.00001 frequency are not reliably genotyped

Across all the variants evaluated for analytical validity using combined cluster plots (n unique=4,585, see Methods), we categorised 27% as high quality (average score>4), 64% as low quality likely false positives (average score<2.5), and 9% as unclear (**Table 1**). There was a strong correlation between the analytical validity quality score and MAF (**Table 1** and **Figure 1**), as well as presence of the variant in either gnomAD^38^ or the 1000 genomes project^39^. For low versus high quality variants, a nonparametric regression analysis estimated the area under the ROC curve to be 0.95 (95% CI = 0.943-0.956); the false positive rate (FPR) at MAF>0.00005 was ~20%, while FPR~60% at MAF>0.00001.

**Figure 1.**
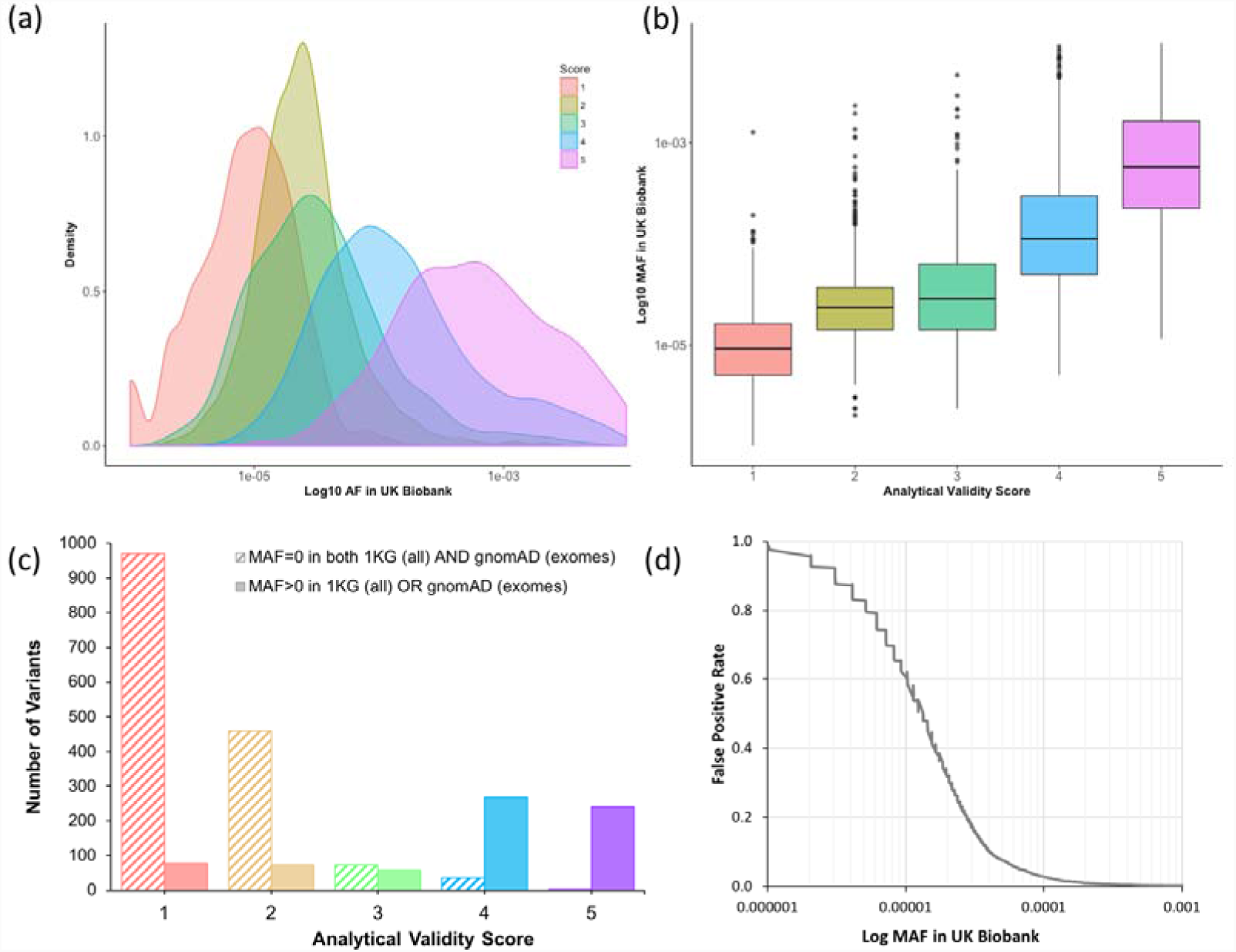
Correlation between MAF and analytical validity quality score. **(a)** Density plot and **(b)** boxplot of manual quality scores (from 1-5, see Supplementary Figure 1) of genotype data in UKB versus minor allele frequency (MAF) for 4,585 putatively clinically important variants, where MAF<0.01, HWE>0.05 and missingness<0.01; **(c)** Histogram of the number of variants at each quality score versus presence or absence of the variant in gnomAD (exome data) or the 1000 genomes project; **(d)** Estimation of the false positive rate (FPR) versus MAF for variants assayed using the UKB genotyping arrays, calculated by grouping quality scores into low (score=1 or 2) and high (score=4 or 5) and using the rocreg command in Stata to fit a ROC curve. Red=score 1; gold=score 2; green = score 3; blue = score 4; purple = score 5.

### Reduced penetrance estimates for known pathogenic variants

The 1,244 high-quality putative pathogenic rare variants, with their ClinVar-associated disease and the allele frequencies in UKB and gnomAD, are shown in **Supplementary Table 2**. Of these variants only 27 were associated with one of the 401 traits we tested against in UKB with p<1×10^-7^ (**Table 2**). Of these, 13 have previously been linked with a dominant disease. For two variants, where penetrance had previously been estimated from large clinical cohorts^40,41^, we found substantially reduced penetrance in our population-based study. Specifically, we observed well-established associations between variants in *PALB2*^40^ and *HOXB13*^41^ and breast cancer and prostate cancer respectively, where the odds ratios were around half the previous estimates from family-based disease studies (in both cases, ~4.5 in UKB versus ~9.5 in the family-based studies)^40,41^. The other 11 variants were causally linked to disease, but penetrance estimates were not available from the literature for comparison. However, we observed that these variants were associated with a related trait in our population-based cohort (**Table 2**), suggesting reduced penetrance versus their presumed monogenic forms. Two PTVs in *FLG* that cause ichthyosis vulgaris^42^ were associated with a 2-fold increased odds of Eczema. A PTV in *TSHR* that causes nonautoimmune hyperthyroidism^43^ was associated with a 3-fold increased odds of hypothyroidism. A nonsynonymous variant in *LRRK2* that causes Parkinson’s disease^44^ was associated with a 5-fold odds of having a parent with Parkinson’s disease. A nonsynonymous variant in *PER3* previously classified as pathogenic for advanced sleep phase syndrome had an odds ratio of only 1.38 for being a morning person^45,46^ compared to a reported 2 hour shift in midpoint sleep. Height, skeletal weight and male pattern baldness were negatively associated with two nonsynonymous variants in *AR* that cause partial androgen insensitivity syndrome^47^. Finally, a nonsynonymous variant in *MYH7*, which has been classified by a ClinGen Expert Panel as pathogenic for hypertrophic cardiomyopathy^48^ was associated with a reduced pulse rate of 5 beats per minute.

We specifically investigated known pathogenic variants and PTVs in MODY genes, where we found two rare variants that were high quality, definitely pathogenic and strongly associated with diabetes (**Table 2**): a very rare stop-gain variant in *GCK* (OR=68 95% CI: 14, 328, P=2×10^-8^), and a nonsynonymous variant (p.R114W) in *HNF4A* (OR=2.9 95% CI: 1.7, 5.0, P=3×10^-4^). Both associated with diabetes in UKB, in-line with previous findings^49-51^. However, the penetrance of the *HNF4A* variant was previously estimated to be up to 75% at age 40-years based on a large MODY diabetes cohort^49^, while we estimate the minimum penetrance to be ~10% from UKB (**Figure 2**). This has important implications for the attributable risk associated with the variant in different cohorts, and the interpretation of genetic test results: if the R114W variant was found in an affected individual following clinical testing, it may still be the primary cause of their diabetes, while incidental discovery of the variant in an unaffected individual would not be predictive.

**Figure 2.**
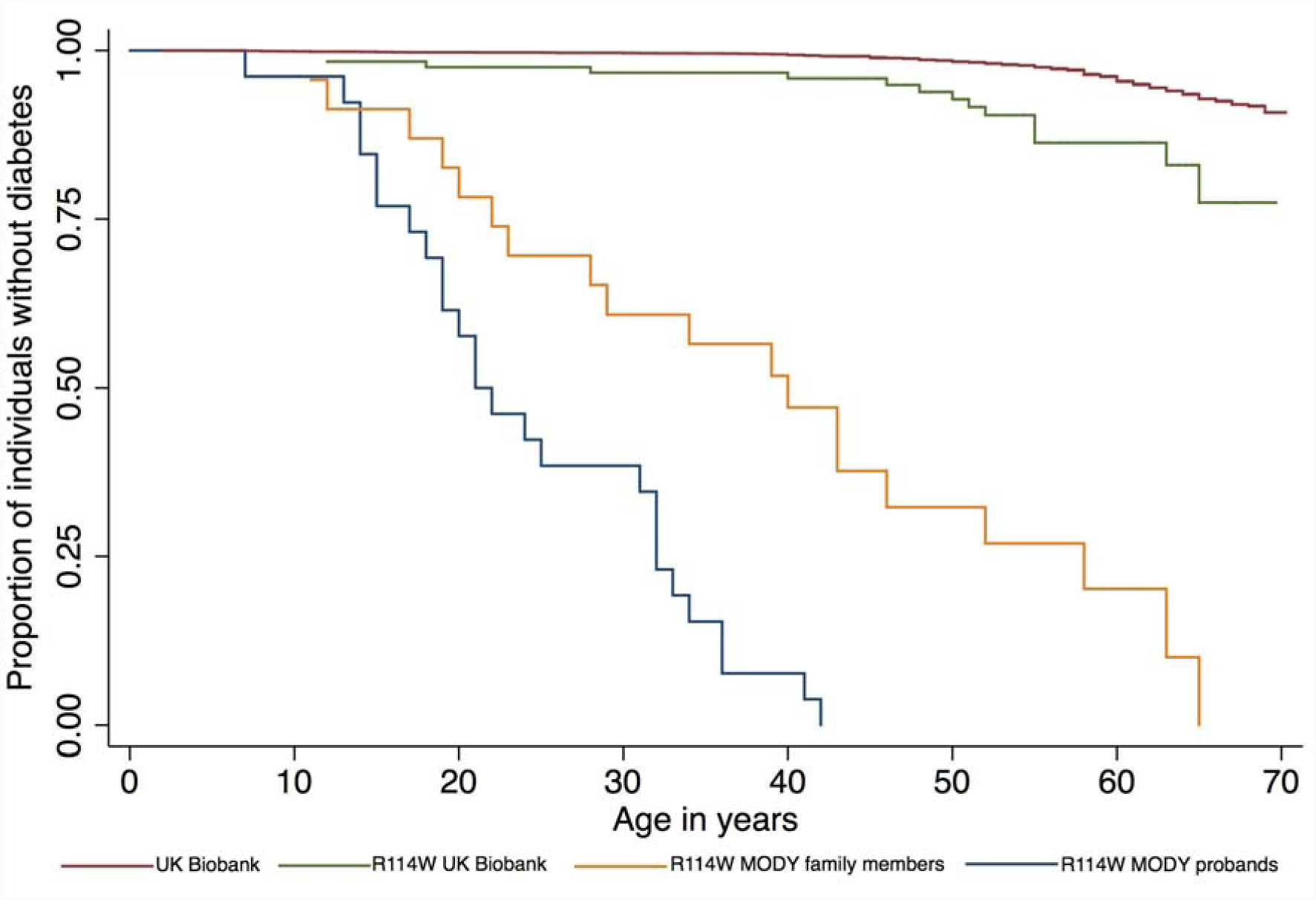
Penetrance estimate for *HNF4A* p.R114W in UK Biobank compared to previously published estimates from MODY cohort studies. A Kaplan-Meier plot of proportion of individuals that are diabetes-free against age for 388,174 individuals from UK Biobank (red line); 122 UK Biobank individuals that are heterozygous for *HNF4A* p.R114W (green line) and 26 MODY referral probands (blue line) and 24 family members of the probands (yellow line) from Laver *et al.*^49^

### Related mild heterozygous phenotypes in autosomal recessive disorders

Of our 27 high-quality, rare putatively pathogenic variants associated with a trait in UKB, 16 have previously been linked with a recessive disease (**Table 2**). We observed associations with milder or related traits in the heterozygous carriers of these monogenic recessive diseases in our population cohort. A nonsynonymous variant in *ERCC4*, which causes recessive xeroderma pigmentosum^52^, and two nonsynonymous variants in *OCA*, which causes oculocutaneous albinism^53,54^ were associated with ease of sunburn. A stopgain variant in *TACR3*, which causes recessive hypogonadotropic hypogonadism^55,56^ was associated with an 8 month increase in age at menarche. In addition, variants in six genes known to cause different recessive blood-related disorders were associated with decreased mean corpuscular volume and/or increased red blood cell distribution width (including *HBB* which causes β-thalassemia but where the carrier state is already known to cause the much milder β-thalassemia minor^57^).

### Refuting previous disease associations

We focused our clinical analysis of variants in DD genes on just PTVs, of which six (including two variants in one gene) were of high-quality and in genes that are reported to cause disease via a haploinsufficiency mechanism (**Table 3**). None of these variants were associated with developmentally relevant traits in UKB (p>0.1), suggesting they are all benign. For three variants, the location of the variant in the gene is notably different from that of known pathogenic variants. *GNAS* is the only one of the five genes with a high probability of being loss-of-function intolerant (pLI)^38^ based on the frequency of loss-of-function variants in ExAC^38^. The stop-gain variant in *GNAS* is present in the highly variable first exon of the gene and is likely to result in nonsense-mediated RNA decay; in contrast, pathogenic variants in *GNAS* that cause Albright hereditary osteodystrophy are located in later, highly constrained exons^58^. Similarly, the stop-gain variant in *TGIF1* is located in the first exon of the gene, where multiple PTVs in gnomAD^38^ are also located, while *TGIF1* pathogenic variants causing holoprosencephaly are located in the final exons of the gene where they affect DNA binding affinity^59^. Finally, a frameshift deletion in *HIST1H1E* is located near the start of the single exon of this gene; however, in contrast, pathogenic *HIST1H1E* frameshift deletions that cause child overgrowth and intellectual disability are located near the end of the exon, where they result in a truncated histone protein with lower net charge that is less effective at binding DNA^60^. Hence, we believe that these three rare PTVs are benign due to their location, despite being PTVs in genes that cause dominant DD via haploinsufficiency.

For the other three variants, our findings are not consistent with the genes causing a dominant DD via haploinsufficiency. First, there was no association between a frameshift variant in the middle of *COL4A3 –* where pathogenic variants are thought to cause a rare dominant form of Alport syndrome (as well as benign familial hematuria)^61,62^ – and albumin creatinine ratios, hearing or any of the development traits in UKB. Similarly, there was no association between either stop-gain or frameshift variants in *RNF135* – where haploinsufficiency is thought to cause macrocephaly, macrosomia and facial dysmorphism syndrome^63^ – with any development traits in UKB. In both cases, given the high-quality genotyping of these variants in UKB and a lack of association with any clinically relevant traits, coupled with a pLI of zero for both genes, the age of the original publications and the lack of enrichment of *de novo* mutations within the DDD study^33^, we suggest that haploinsufficiency in these genes is not a cause of a severe DD.

## CONCLUSIONS

Previous studies have been unable to analyse rare variants in sufficiently large population-based studies to establish pathogenicity and lower-bounds for penetrance. Large population cohorts such as UKB provide an opportunity to investigate the relationship between genes and disease. However, the absence of genome-wide sequencing data has thus far minimised the impact of UKB in the rare disease community. We have established a method for evaluating the analytical validity of rare variants genotyped by microarray, using combined intensity plots for individual variants across all genotyping batches. Although we initially tried to examine variant cluster plots for each batch separately, as recommended by UKB, this proved impossible due to the rarity of most clinically important variants. MAF was an extremely good predictor of the likelihood of a variant being genotyped well by the UKB arrays (**Figure 1**). At MAF>0.00005 (~50 heterozygous individuals) FPR~7% and most variants were well genotyped, while FPR~60% at MAF>0.00001 (~10 heterozygous individuals), and we classified all variants at MAF< 0.000005 (~5 heterozygous individuals) as being low quality. This has important implications for epidemiological research carried out uncritically using these data. Although many rare variants in UKB are well genotyped with the arrays, the rarer the variant, the more likely it is to be poor quality and therefore yield false associations.

A limitation of our work is that we did not attempt to confirm the variants using an independent assay. However, most researchers using data from UKB will be similarly unable to attempt independent variant confirmations, and thus a method for evaluating the genotyping quality of rare variants directly from the data has widespread utility. The validity of our method is supported by our ability to replicate numerous previous findings of well-known, clinically important variants classified as pathogenic in ClinVar (**Table 2**, plus additional well-established associations for variants where MAF>0.01). In addition, our analyses of likely pathogenic variants in two disease subtypes (MODY and DD) were independent of any potential biases or misclassification errors associated with ClinVar, and the findings were consistent with our prior expectations. We expected there to be a small number of individuals in UKB with monogenic subtypes of diabetes and we found two pathogenic variants that associated with appropriate traits in UKB (**Table 2**) and were thus able to lower the previous penetrance estimate for a pathogenic variant in *HNF4A* (**Figure 2**). In contrast, we did not expect there to be any instances of severe DD, due to the rarity of the condition, the relatively senior age of the UKB population and the inherent challenges of consenting individuals with severe DD to population biobanks^64^. We are therefore confident that the PTV variants identified in dominant DD genes in UKB are benign (**Table 3**), and in refuting previous associations between haploinsufficiency in *RNF135* and *COL4A3* and dominant DD (which has no bearing on the asserted relationship between the latter and either recessive DD or alternative mechanisms of disease).

In this study, we have shown that population genetic data can be used to estimate lower bounds for the penetrance of pathogenic disease-causing variants, and refine our understanding of the links between rare variants (MAF<0.01) and monogenic diseases. Performing a similar analysis on ultra-rare variants (MAF<0.00001) will require large-scale sequencing data rather than genotyping arrays. Although population-based studies will be biased in the opposite direction from clinical studies, i.e. towards healthy individuals, they are nonetheless crucial for interpreting incidental or secondary findings from clinical testing, and for informing direct-to-consumer genetic testing. At this point, we are left with some fundamental conceptual questions about the nature of “monogenic” disease. When should variants exhibiting reduced penetrance – a term frequently used in the diagnosis of rare genetic disease – be called risk or susceptibility factors – terms generally used in the study of common disease? When should a gene-disease relationship be termed variable expressivity rather than normal variation? Should “pathogenic” be reserved only for highly penetrant variants that cause a tightly defined disease entity, or can it apply to any variant associated, however weakly, with a clinically-relevant phenotype? As genome-wide sequencing becomes widely used in routine clinical practice, research cohorts and direct-to-consumer testing, understanding this spectrum will become both increasingly important and tractable.

## ACKNOWLEDGEMENTS

This research has been conducted using the UK Biobank Resource. This work was carried out under UK Biobank project number 871.

## TABLES

**Table 1. Evaluated variants.** Number of variants manually evaluated for analytical validity in different MAF bins, with quality scores grouped into false positive (FP, score=1 or 2), unclear (score=3) and true positive (TP, score=4 or 5).

**Table 2. Pathogenic variants.** Reduced penetrance, variable expressivity and carrier phenotypes for rare (MAF<0.01) ClinVar pathogenic variants with genome-wide significant associations in UKB.

**Table 3. Benign variants.** Classification of likely pathogenic variants in maturity-onset diabetes of the young (MODY) and developmental disorders (DD) from UKB.

**Supplementary Table 1. Curated traits included from UKB.**

**Supplementary Table 2. 1,244 high quality putative pathogenic variants analysed.**

## SUPPLENTARY FIGURES

**Supplementary Figure 1.**
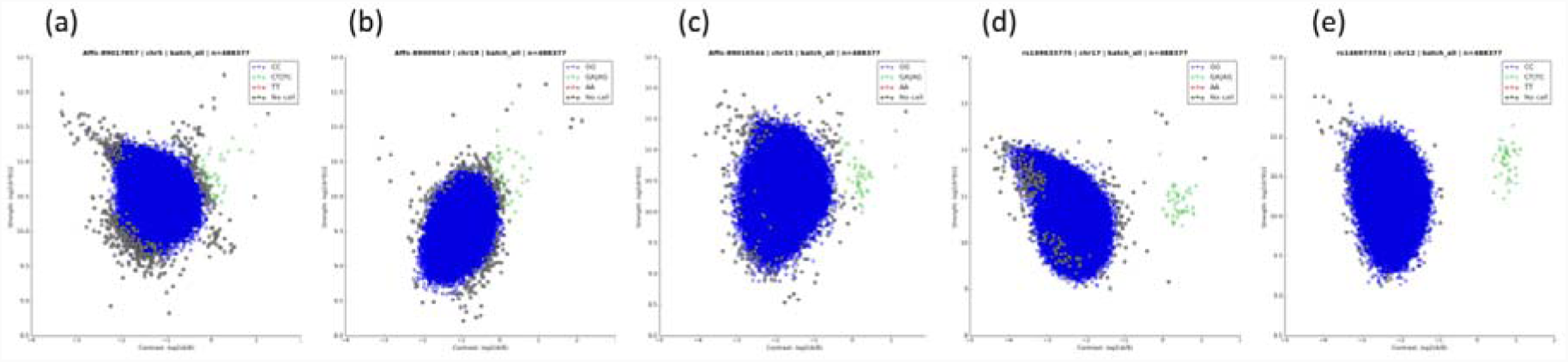
Combined cluster intensity plots. Intensity plots combined across all batches are shown for five variants, all with a UKB MAF = 0.00004. The clustering quality of heterozygous variants was manually assessed and ranked from 1-5. **(a)** Score 1 = poor quality, no discernible separate clusters; **(b)** Score 2 = poor quality, no discernible separate clusters but noisy data; **(c)** Score 3 = unclear/uncertain; **(d)** Score 4 = good quality, clearly separable clusters but noisy data; **(e)** Score 5 = good quality, clear separation between clusters.

**Figure 2.**
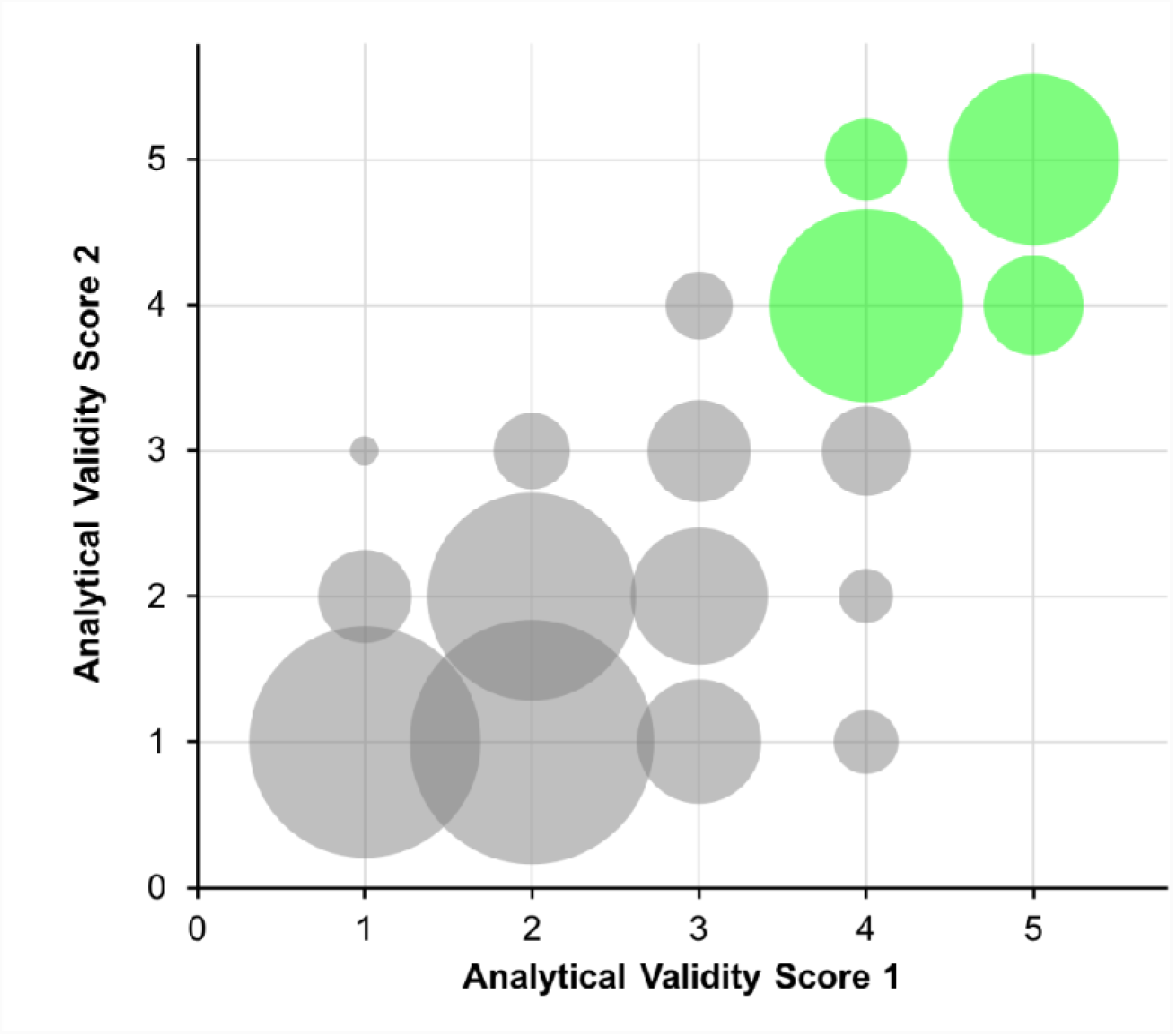
Comparison of quality scores between two independent scorers. Two scientists independently scored the quality (from 1-5, see Figure 1) of combined cluster plots for 750 variants. The R^2^ between their scores was 0.8, and there was a 95% agreement in low quality (score=1 or 2) versus high quality (score=4 or 5) variants.

